# The stabilized gaze signalling hypothesis and the evolution of human eye appearance

**DOI:** 10.1101/2025.10.16.682340

**Authors:** Fumihiro Kano, Hiromi Kobayashi, Kazuhide Hashiya

## Abstract

Human eyes are commonly described as having a ‘white sclera’: they typically combine scleral exposure with depigmentation, making eye direction readily visible. However, variation in scleral pigmentation among other primates has challenged claims of human uniqueness and renewed debate about its role in gaze communication. Here we develop the stabilized gaze-signalling hypothesis (SGSH), which makes four predictions: (1) depigmented, exposed sclera supports more robust eye-gaze direction discrimination than pigmented sclera across variable viewing conditions by preserving joint visibility of the iris and eye outline; (2) coordination benefits increase when viewing conditions vary or monitoring more group members increases attentional demands, relative to sustained close-range dyadic interaction; (3) humans show greater receiver sensitivity to eye-gaze cues than other primates across development; and (4) heritable morphological variation was available for selection, with signalling benefits potentially exceeding costs of revealing information. Existing evidence supports perceptual benefits under some conditions; the other predictions require further testing. We also consider selective pressures, alternative origins, and complementary functions. The SGSH provides a revised framework for testing whether social benefits from maintaining readable eye direction during group activities could explain the spread and maintenance of depigmentation of exposed sclera in humans.

## Introduction

Human eyes are striking in their appearance. Compared with other primates, human eyes typically combine a widely exposed sclera with depigmentation. In comparative morphological analyses, Kobayashi and Kohshima (1997, 2001) described this configuration as uniquely human and proposed the gaze-signalling hypothesis (GSH): white sclera evolved to make gaze direction conspicuous for communication. Tomasello et al. (2007) extended this account with the cooperative eye hypothesis (CEH): conspicuous eyes evolved to facilitate mutually beneficial coordination through joint attention and communication among partners unlikely to exploit gaze information to the looker’s disadvantage.

Comparative research has since shown considerable variation in scleral pigmentation, exposure, and contrast, both within species and among primates (Caspar et al., 2021; Clark et al., 2023; Kano et al., 2022a; Mayhew and Gómez, 2015; Mearing et al., 2022; Perea-García et al., 2019, 2022, 2024). Humans are not categorically distinct in each of these traits. Perea-García et al. (2025) interpreted these findings, together with limitations in the behavioural evidence, as grounds for moving beyond the GSH/CEH. They argued that human ocular pigmentation is neither uniquely distinctive nor uniform within the species, and that evidence for its contribution to eye-gaze signalling remains insufficient to support the proposed adaptive explanation. They therefore called for greater attention to alternative explanations of ocular appearance, including photoregulation, sexual selection, and non-referential social signalling.

These findings challenge categorical claims of human uniqueness, but do not by themselves rule out quantitative differences between humans and other primates or a gaze-signalling explanation of human eye appearance. Kano (2023) distinguishes the visibility of eye position within the face from the visibility of eye-gaze direction. For the latter, the iris must be visible relative to the eye outline. The proposed advantage of uniformly white sclera is that it helps maintain this visibility across changes in viewing conditions. Building on these arguments, we propose the stabilized gaze-signalling hypothesis (SGSH) as a refinement of the GSH and CEH. It retains their core gaze-communication account while developing Kano’s (2023) arguments into linked predictions with explicit tests and falsification criteria.

The SGSH proposes that depigmentation of exposed sclera evolved to stabilize eye-gaze direction discrimination across the variable viewing conditions encountered in human social life. Here, variation refers to the range of conditions under which eyes are viewed: people may be seen at different distances, under different illumination, away from the centre of vision, or only briefly. The proposed perceptual advantage concerns locating the iris within a visible eye outline. Maintaining the visibility of this spatial relationship across these conditions could make others’ eye-gaze directions easier to assess, particularly in large groups where members are spatially distributed and viewed under different conditions, or where monitoring more individuals increases attentional demands. We further propose that the use of such information in group activities provided opportunities for selection on eye morphology and that enhanced receiver sensitivity to eye-direction cues also evolved.

Table 1 defines the terms used throughout this review, and Table 2 states four linked predictions concerning the robustness of eye-gaze direction discrimination (P1), benefits for human group activities (P2), quantitative receiver specialisation (P3), and heritable variation permitting selection where benefits exceed costs (P4). P1, P2 and P4 are necessary premises of the SGSH, while P3 is a prediction derived from the possibility that signals and receiver sensitivity coevolved (Dawkins and Krebs, 1978). For each prediction, we distinguish existing evidence from open questions and identify results that would challenge the account. Research on P1 is comparatively advanced, although the range of conditions tested remains limited. Direct tests of P2–P4 are much scarcer, and these components remain working hypotheses requiring further research. In this respect, we agree with Perea-García et al. (2025) that the available evidence does not yet establish an adaptive explanation of human eye appearance. The following sections assess the evidence and open questions for each prediction; Table 2 identifies tests and results that would challenge these predictions.

**Table 1.** Key terms and distinctions used in this review.

| **Term** | **Definition and scope** |
|---|---|
| Eye-gaze signalling | Conveying information about eye direction through the appearance or movement of the eyes. Eye-gaze cues specify eye direction, whereas head-gaze cues specify head orientation; gaze-direction discrimination can be based on both cue types. |
| Scleral exposure and pigmentation / depigmentation | Scleral exposure is the extent of the scleral region visible within the eye opening. Here, scleral pigmentation refers to the visible colouration of this region, including pigmentation in the overlying conjunctiva; depigmentation denotes reduced or absent pigmentation. These terms describe external appearance rather than the pigmentation of scleral tissue itself. |
| Viewing conditions and attentional demands | Viewing conditions are the circumstances under which eyes are observed, including distance, illuminance, retinal eccentricity, and viewing duration. Tests should sample conditions representative of relevant everyday interactions. Attentional demands concern allocating attention across individuals or cues; they can interact with viewing conditions. |
| Co-visibility of the iris and eye outline | The eye outline is the visible boundary of the eye opening and provides a spatial reference for the iris. When the iris is light-coloured, the pupil may instead provide the positional cue. Co-visibility means that the iris and this outline are simultaneously discernible under the same viewing conditions. Maintaining co-visibility is one proposed mechanism supporting robust eye-gaze direction discrimination. |
| Eye-gaze direction discrimination | An observer’s ability and efficiency in perceptually distinguishing eye-gaze directions. |
| Robustness of eye-gaze direction discrimination | The extent to which eye-gaze direction discrimination is maintained across changes in viewing conditions. |
| Receiver sensitivity | An observer's ability to discriminate eye-gaze direction and propensity to attend to and rely on eye-gaze cues in guiding attention and behaviour. Perceptual discrimination, spontaneous cue use, and relative cue weighting are related but distinct components. |
| Behavioural measures of gaze perception and cue use | Discrimination tasks assess accuracy, estimation error, response time or thresholds, often after training in nonhuman animals. Cue-use measures include gaze following (overt orienting), gaze cueing (faster or more accurate responses to gaze-congruent targets), and object choice. Spontaneous cue use refers to responses without explicit instruction or training to use the gaze cue. Relative eye–head cue weighting is the relative influence of these cues on gaze perception or behaviour within a specified task. |

**Table 2.** Predictions of the SGSH and approaches to testing them.

| **Component** | **Prediction** | **Tests and measurements** | **Contrary results and constraints** |
|---|---|---|---|
| P1. Robustness of eye-gaze direction discrimination | Human-like eyes, combining scleral exposure with depigmentation, maintain eye-gaze direction discrimination across a broader, ecologically relevant range of viewing conditions than eyes with pigmented sclerae. | • Key viewing conditions, such as distance, illuminance, retinal eccentricity, and viewing duration, can be varied across a diverse range representative of everyday interactions.<br>• Discrimination performance can be assessed from accuracy and response time, considered together to account for speed–accuracy trade-offs.<br>• Image-based measures of iris–eye outline co-visibility can characterise the visual information available under this proposed mechanism; their relationship to behavioural discrimination requires testing. | • No robustness advantage across diverse everyday conditions, or benefits confined to a narrow set of experimental conditions. |
| P2. Benefits for human group activities | More robust eye-gaze direction discrimination supports social coordination. Compared with sustained, close-range dyadic interaction, human-like eyes are predicted to confer greater benefits when spatial dispersion creates variable viewing conditions or when monitoring more group members increases attentional demands. | • Screen-based tasks can compare eye types in a factorial design varying the number of faces and viewing conditions to test their separate effects and interaction.<br>• Live tasks can test eye-type effects on coordination and performance during natural interactions in dyads and larger groups, where viewing conditions, spatial arrangement, and attentional demands vary, while accounting for head direction, speech, and gestures. | • Screen-based tests: no reliable advantage of human-like eyes, or no increase in their benefit as the number of faces or viewing conditions vary. Benefits limited to a narrow set of conditions would restrict the prediction's scope.<br>• Live tasks: no coordination or performance benefit, or no greater benefit than in sustained, close-range dyadic interaction. Other cues may fully compensate without additional time or effort. |
| P3. Receiver specialisation | Compared with other primates, humans show greater receiver sensitivity to eye-gaze cues across development. | • Crossed observer–stimulus species designs can compare eye-gaze direction discrimination, spontaneous eye-cue use, and relative eye–head cue weighting across development. Head-cue responses provide a within-observer reference when motivation and training complicate comparisons.<br>• Comparisons should account for visual acuity, stimulus-species experience, training, and motivation. | • Equivalent or greater receiver sensitivity in nonhuman primates in matched discrimination and cue-use comparisons across development. |
| P4. Heritable variation and selection | Heritable variation in eye morphology was available when selection acted, and the benefits of eye-gaze signalling could exceed the costs of revealing information. | • Heritability of ocular pigmentation can be estimated by combining standardised measurements with estimates of relatedness from family records or genomic data, while accounting for age and shared environmental effects.<br>• The spread and maintenance of depigmentation can be modelled using empirically grounded estimates of signalling benefits and information-leakage costs, together with relevant ecological and social parameters. | • Evidence ruling out appreciable additive genetic variation would limit the potential for an evolutionary response to ongoing selection in the population studied, without excluding past selection.<br>• Failure of models across empirically plausible benefits, costs, and ecological and social conditions to support the spread or maintenance of depigmentation would weaken the proposed selection scenario. |

## Prediction 1 Robustness of eye-gaze direction discrimination

P1 is that human-like eyes, combining scleral exposure with depigmentation, maintain eye-gaze direction discrimination across a broader, ecologically relevant range of viewing conditions than eyes with pigmented sclerae (Table 2). Key examples include viewing distance, illuminance, retinal eccentricity, and viewing duration, all of which vary naturally in everyday human group interactions.

### Spatial information for eye-gaze direction discrimination

Head orientation and eye direction provide related but distinguishable information. Head orientation can be estimated from the nose, mouth, facial contour, and overall alignment even when the eyes are poorly resolved; eye-gaze direction can specify where within that broader visual field the individual is looking (Emery, 2000; Kobayashi and Hashiya, 2011; Kobayashi and Kohshima, 2001; Tomasello et al., 2007). For the spatial mechanism emphasised here, both the iris and the eye outline that provides a reference for its position must remain discernible (Kano, 2023; Kano et al., 2022a, 2022b). Detecting either feature alone is insufficient to determine their spatial relationship. Figure 1 illustrates this relationship. Facial features can continue to indicate head orientation when the eye outline is indistinct, whereas locating the iris within the eye opening requires both features to remain visible.

**Figure 1.**
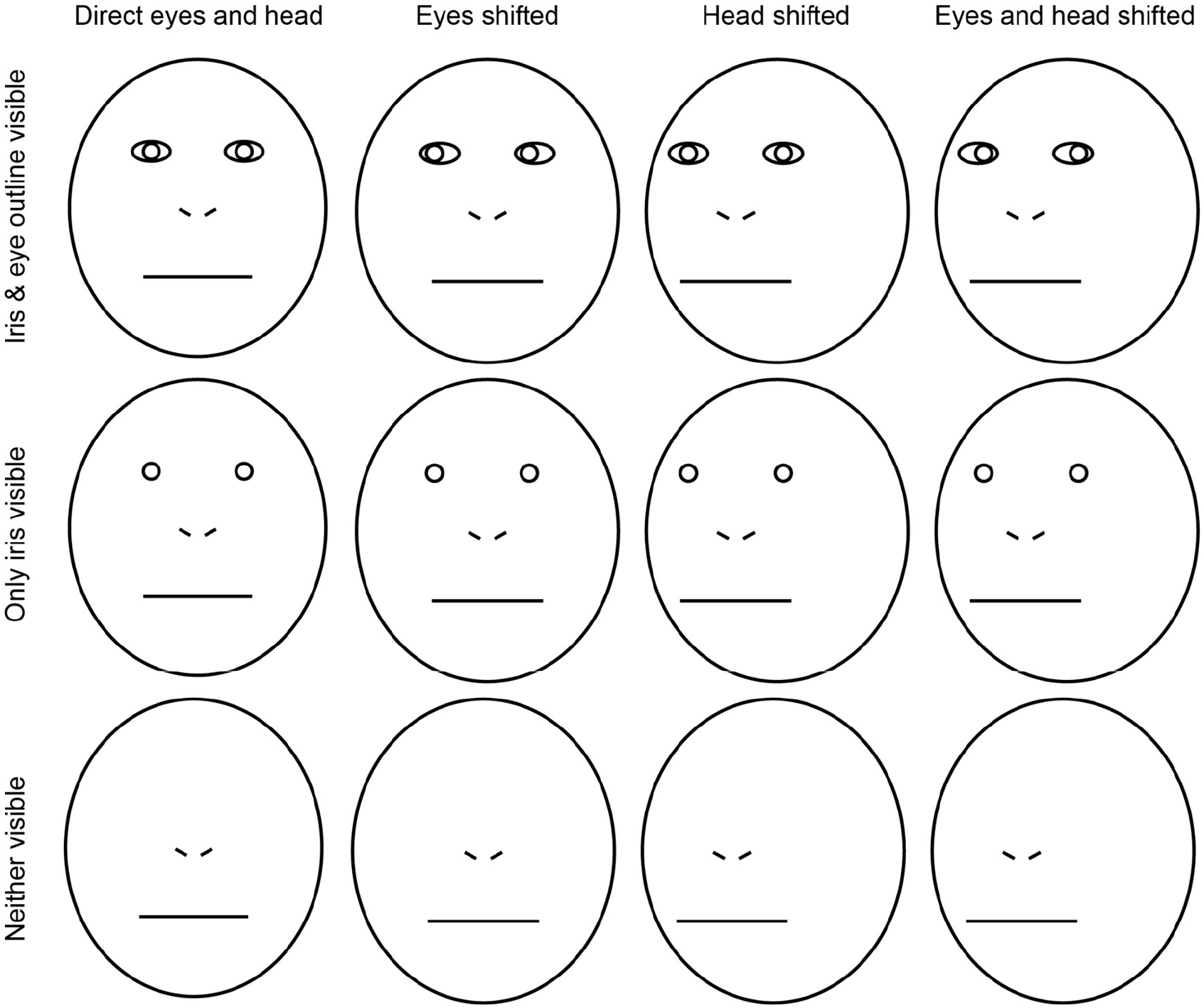
Schematic separation of eye-gaze and head-orientation cues. The columns show direct eyes and head, eyes shifted within the eye openings, head shifted with the irises centred within the eye openings, and both eyes and head shifted. Head shifts are depicted by translating the eye openings, irises, nose and mouth together relative to the outer face contour; eye shifts displace the irises relative to the eye openings. Head and eye shifts are 15% of the maximum face width and maximum eye-opening width, respectively. The rows preserve the corresponding feature positions while showing both iris and eye outline (top), only the iris (middle), or neither (bottom). The top row provides a visible reference for iris position within the eye opening.

Figure 3A illustrates alternative colour and luminance arrangements that can support co-visibility of the iris and eye outline. Examples include a dark iris against a light sclera, as typically seen in humans, and a light iris against a dark sclera, as typically seen in chimpanzees and mountain gorillas (Kano et al., 2022a). Both configurations can provide directional information, provided that the eye outline remains discernible from the surrounding skin or hair. The four image-mask pairs capture these different contrast configurations without privileging the human-typical polarity. Regional colour differences alone do not capture all the visual information relevant to eye-gaze direction discrimination. Eyelid margins, eyelashes, and local shading can delineate the eye outline even when the sclera and surrounding skin have similar average colouration (Kano et al., 2022a). Eye-gaze direction discrimination also relies strongly on mid-to-high spatial frequencies, which convey relatively fine visual detail (Vida and Maurer, 2015). Measures of eye appearance should therefore consider local edges as well as regional colour differences. The following sections clarify which of these features existing approaches assess.

### Scleral exposure

In their original comparisons, Kobayashi and Kohshima (1997, 2001) highlighted the limited exposure of the sclera in many nonhuman primates relative to humans. Kobayashi and Kohshima (2001) reported that the proportion of scanning movements made by the eyes alone increased with scleral exposure across the primates they observed, and related this pattern to body size. In species that shift their gaze predominantly through head movements, head orientation may provide useful directional information even when little sclera is exposed. The contribution of scleral colouration to eye-gaze direction discrimination is therefore best considered in relation to how much sclera is visible and how the eyes move within the eye opening.

Kobayashi and Kohshima (2001) quantified scleral exposure using the Sclera Size Index (SSI), the ratio of eye-opening (visible eyeball region) width to iris diameter. Kano et al. (2022a) reassessed human distinctiveness using the Sclera Area Index (SAI), the proportion of the eye-opening area occupied by sclera, and found overlap between humans and other great apes. This finding challenges an explanation based simply on greater scleral exposure in humans and shifts attention to how exposure and colouration jointly support the co-visibility of the iris and eye outline. Within-human comparisons likewise caution against interpreting differences in SSI as differences in exposed scleral area. In 100 Czech adults, Danel et al. (2020) found higher SSI in men than women, but no significant sex difference in the percentage of the eye-opening area occupied by sclera, measured using their sclera surface ratio (SSR). Boyer-Brosseau et al. (2025) additionally found poor-to-good inter-rater reliability for SSI. Lee et al. (2013) found that observers discriminated gaze direction more accurately in schematic eyes as the vertical eye opening increased from disgust-like narrowing to fear-like widening. This advantage persisted when the eyes were inverted to reduce perceived fear, suggesting that it arose from the physical configuration of the eyes rather than perceived emotion alone. However, widening changes both eye shape and the amounts of iris and sclera visible, so this experiment does not isolate the contribution of scleral exposure or establish whether naturally occurring differences in scleral exposure among individuals have similar effects.

Perea-García et al. (2025) presented the golden langur and southern pig-tailed macaque as examples of nonhuman primates whose scleral (or peri-iridal) brightness overlaps with that of humans, challenging the claim that light scleral appearance is uniquely human. However, evaluating the implications of this overlap for eye-gaze signalling also requires considering how much of the light tissue is visible to an observer. In the illustrated views of these species, little of this tissue is exposed (Figure 2). Figure 2 places these examples alongside human and other great ape eyes drawn from Kano et al. (2022a). These examples serve an explanatory purpose; systematic measurements of exposure and co-visibility, linked to behavioural tests under comparable viewing conditions, are still needed.

**Figure 2.**
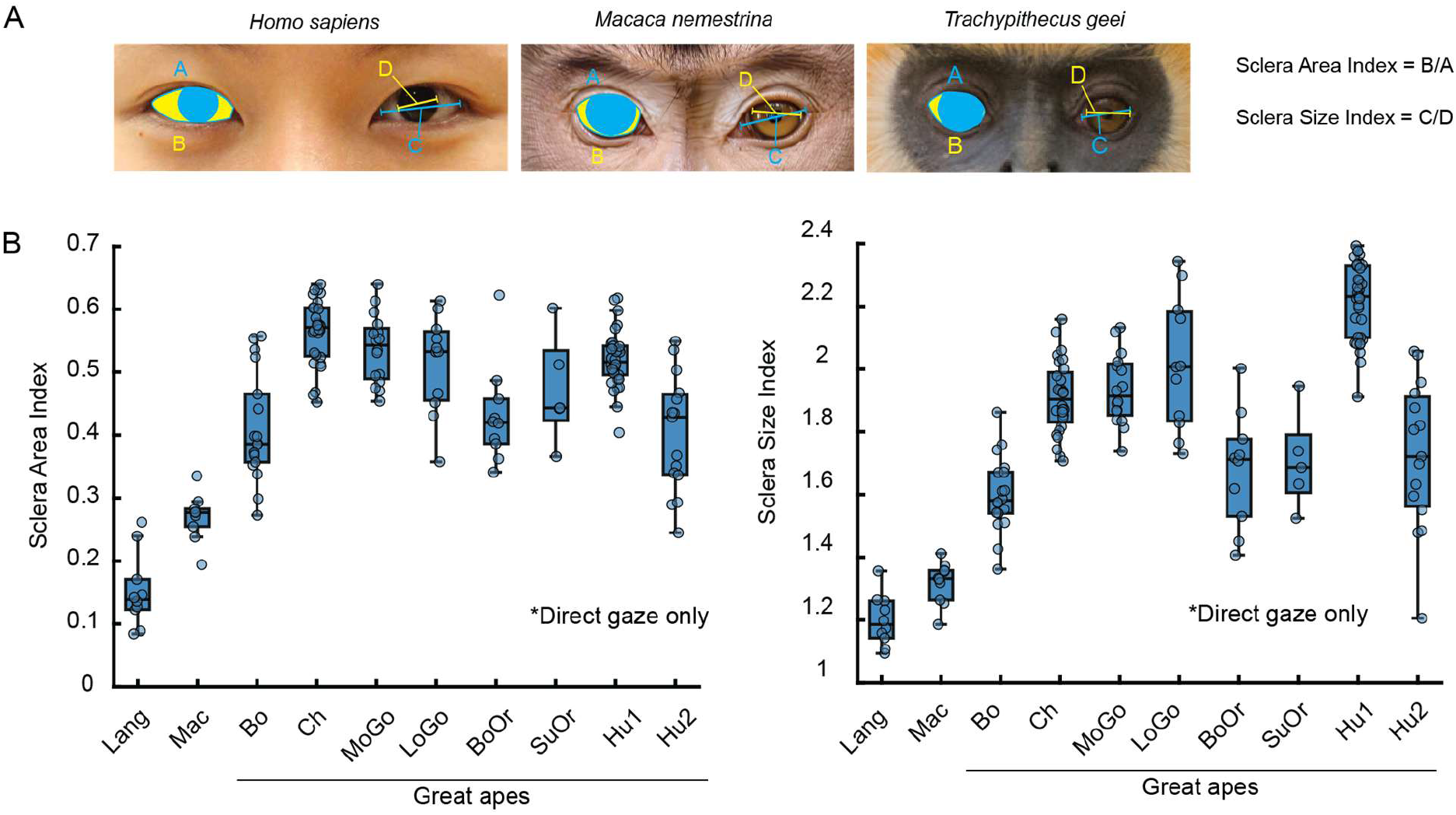
Sclera Area Index (a 2D measure: sclera area divided by eye opening area; Kano et al., 2022a) and Sclera Size Index (a 1D measure: longest dimension of the eye opening divided by the longest dimension of the iris; see also Boyer-Brosseau et al., 2025 for these measures). Points represent individual direct-gaze eye images. For the golden langur and southern pig-tailed macaque, 10 publicly available internet images were sampled for each species. All nonhuman great ape images were sourced from Kano et al. (2022a). Human images were sampled from the Columbia Gaze Dataset (Smith et al., 2013) and Figure 4 of Perea-García et al. (2025). Code and data are shared at Open Science Framework (https://osf.io/z6753/?view_only=5f6f393077194fb789c7fe5c44bbe06e). In this figure, the human image is from the Columbia Gaze Dataset; the macaque image is used under Shutterstock Standard License; the langur image © Yeray Seminario, iNaturalist, licensed under CC BY-NC 4.0 (https://creativecommons.org/licenses/by-nc/4.0/). Lang, golden langur; Mac, southern pig-tailed macaque; Bo, bonobo; Ch, chimpanzee; MoGo, mountain gorilla; LoGo, lowland gorilla; BoOr, Bornean orangutan; SuOr, Sumatran orangutan; Hu1, human (Smith et al., 2013); Hu2, human (Figure 4 of Perea-García et al., 2025). See Supplemental Material for methodological details. This preliminary sample illustrates differences in visible exposure; it does not estimate species-wide distributions or establish functional differences in eye-gaze direction discrimination.

### Variation within and among species

Comparisons among species need to account for individual variation in the visible configuration of the eye. Variation in scleral pigmentation has been reported in gorillas, including individuals with extensive depigmentation (Mayhew and Gómez, 2015), and in chimpanzees, including the Ngogo population (Clark et al., 2023). Both direct and averted gaze should be considered because eye rotation can expose peripheral scleral regions whose colouration is less apparent during direct gaze, as illustrated by Mayhew and Gómez (2015). Variation in contrast polarity also matters: chimpanzees can have a relatively light iris against a dark sclera, reversing the typical human arrangement (Perea-García et al., 2019; Kano et al., 2022a; Whitham et al., 2022a).

The relevant question is whether the iris and the eye outline remain jointly discernible across these configurations. Figure 3 presents a reanalysis of the images from Kano et al. (2022a), using the mask comparisons described above to quantify contrast configurations supporting co-visibility. This approach accommodates different combinations of iris, scleral, and surrounding skin colouration, including reversed contrast polarity, and illustrates differences between human and other great ape eyes in these image-based measures. The maximum ΔE selects the strongest of the sampled contrast configurations, allowing different colour arrangements to contribute to the measure. It is a configuration-level proxy motivated by co-visibility, not an independent measure of iris or eye-outline visibility.

**Figure 3.**
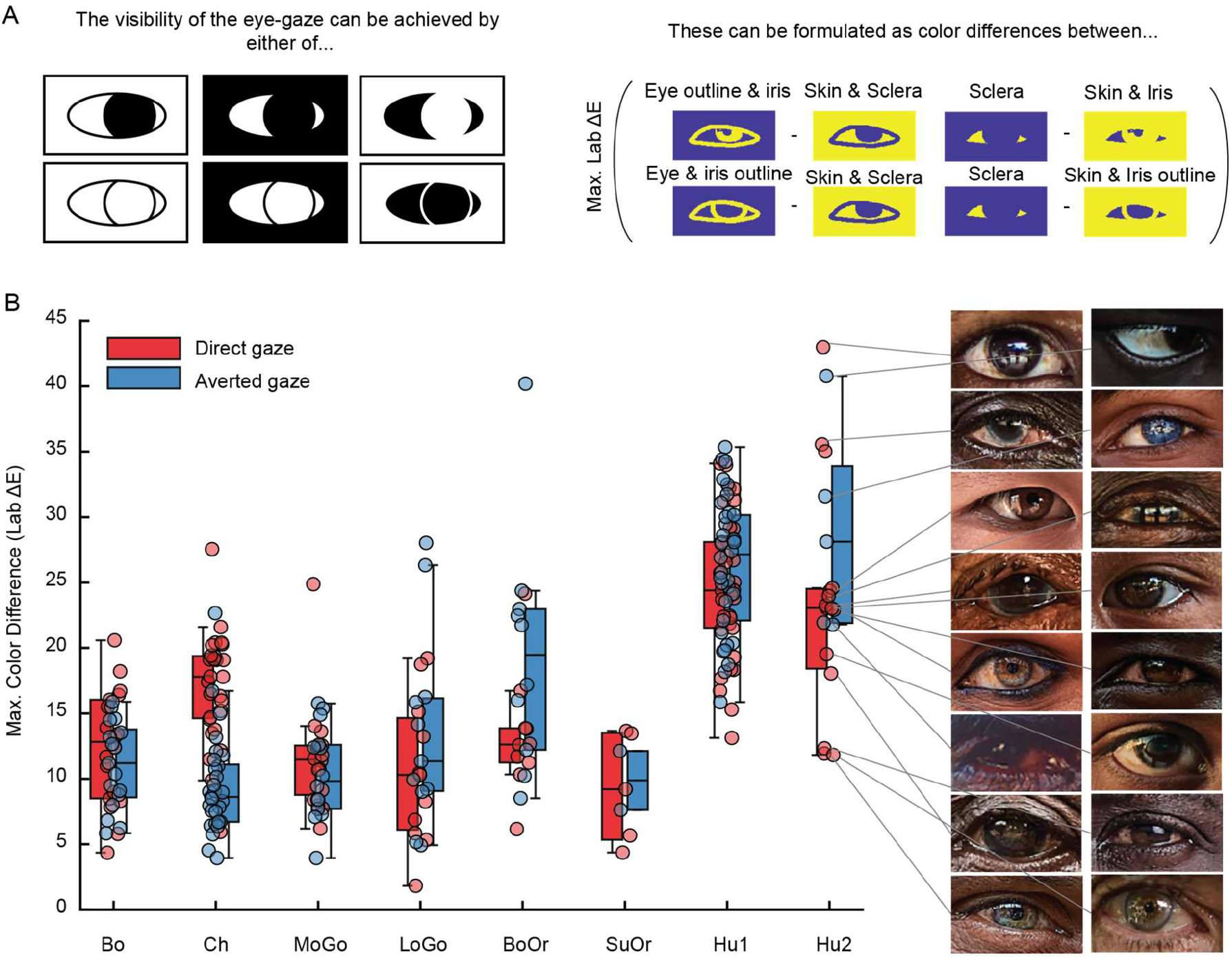
(A) Schematic representation of the colour patterns when eye-gaze is visible and the four image mask pairs used to quantify these patterns. The mask pairs represent alternative spatial colour arrangements in which the iris and eye outline can both be delineated. (B) Colour difference between the mask pairs (CIELAB ΔE). For each image, the maximum ΔE across the four mask pairs is reported as an illustrative measure of gaze-related colour contrast. Each point represents one eye image; red and blue indicate direct and averted gaze, respectively. All images were standardised for tonal range, and colour measurements were conducted following the protocol of Kano et al. (2022a). Code and data are shared at Open Science Framework (https://osf.io/z6753/?view_only=5f6f393077194fb789c7fe5c44bbe06e). Image © Marios Forsos licensed under CC BY-NC-SA 4.0 (https://creativecommons.org/licenses/by-nc-sa/4.0/). Bo, bonobo; Ch, chimpanzee; MoGo, mountain gorilla; LoGo, lowland gorilla; BoOr, Bornean orangutan; SuOr, Sumatran orangutan; Hu1, human (Smith et al., 2013); Hu2, human (Figure 4 of Perea-García et al., 2025). See Supplemental Material for methodological details. These image measurements illustrate the proposed configuration; their relationship to behavioural eye-gaze direction discrimination requires testing.

Human eyes also vary in scleral pigmentation, as emphasised by Perea-García et al. (2025). However, the maximum ΔE values for the partially pigmented human examples highlighted in that study (Hu2 in Figure 3B) show substantial overlap with those for the human images from the Columbia Gaze Dataset (Smith et al., 2013) analysed by Kano et al. (2022a) and reanalysed here (Hu1), although some Hu2 values extend beyond the Hu1 range. This preliminary comparison illustrates how such variation can be evaluated using the same measures. A systematic assessment using a more broadly representative sample of human variation remains necessary.

### Effects of viewing conditions

Viewing distance, illuminance, retinal eccentricity, and viewing duration vary naturally in everyday human group life as individuals move, enter shade, shift attention, and glance briefly at one another. These variations affect the spatial and contrast information available to observers. The robustness of eye-gaze information therefore needs to be assessed across these viewing conditions.

Kano et al. (2022a) examined how darkening and blur affect the joint visibility of the iris and eye outline in images of humans and other great apes. Darkening left iris–sclera colour differences discernible in most species under an approximate human colour-difference criterion, whereas colour differences defining the eye outline became barely detectable in nonhuman apes; human images retained both contrasts. In a separate analysis, increasing Gaussian blur reduced the proportion of iris- and eye-outline edges recovered by edge detection, with human images retaining more of both at the strongest blur. These results suggest that uniformly white sclera helps preserve both the iris and its surrounding reference boundary when image contrast or spatial detail is reduced.

Whitham et al. (2022a, 2022b) incorporated receiver visual characteristics and viewing distance into models of nonhuman primate eyes. A particular strength of this approach is its simulation of species-specific vision, allowing eye appearance to be evaluated in relation to the visual capacities of relevant receivers. Their models quantified chromatic and achromatic discriminability among eye and surrounding facial regions, showing that detectable regional contrasts can occur without human-like white sclera. However, they did not directly compare human and nonhuman eye configurations. Although distance-related blurring was incorporated, the reported regional contrasts do not directly quantify the edge information defining the eye outline.

Overall, image-based approaches can analyse key visual features for eye-gaze direction discrimination, such as regional colour differences and iris and eye-outline edges. These analyses provide proxy measures that can help identify informative conditions for subsequent behavioural tests. They also allow systematic sampling across multiple levels within an ecologically relevant range, revealing gradual declines, thresholds, and changes in the relative advantage of different eye configurations that comparisons between two extremes may miss. However, these simulations approximate only selected aspects of natural viewing. Blur can isolate losses of spatial detail associated with distant or peripheral viewing, and darkening can approximate reduced illumination, but changes in retinal image size, viewing angle, peripheral crowding, local shadows, reflections, and visual adaptation also affect what observers see. Interpretation therefore requires standardised images, appropriate modelling of receiver visual characteristics, and ultimately validation against behavioural eye-gaze direction discrimination under relevant viewing conditions.

### Eye-gaze direction discrimination test

Behavioural experiments provide more direct tests of eye-gaze direction discrimination than image-based measures alone. Depending on task demands, both accuracy and response time are informative, and considering them together helps account for possible speed–accuracy trade-offs.

Yorzinski and Miller (2020) manipulated scleral colouration in photographs of human faces and asked human observers to find a direct-gaze face among averted-gaze faces, or vice versa. Search was generally faster for naturally white sclera or sclera lighter than the iris than for sclera matching or darker than the iris. For small faces, sclera darker than the iris were associated with slower manual responses and lower accuracy than naturally white sclera. Yorzinski et al. (2021) extended this approach to simulated nighttime viewing by combining image darkening, a blue colour shift, Gaussian noise, and blur. Naturally white sclera facilitated performance in both daytime and simulated nighttime conditions. These findings support P1 under the tested conditions. However, the use of human observers and human faces leaves open a contribution from perceptual tuning to familiar eye configurations or reliance on their typical luminance relationships (Ricciardelli et al., 2000; Olk et al., 2008). The combined nighttime manipulation also prevents its individual components from being evaluated separately.

Kano et al. (2022b) used a crossed observer–stimulus design, testing humans in a left/front/right eye-gaze direction discrimination task and chimpanzees in a three-face visual-search task, using images of both species. Observers accurately discriminated gaze direction from both eye types in larger, brighter images, while white-sclera eyes yielded higher accuracy when image size and brightness were reduced together. Reversing eye contrast polarity impaired performance for human eyes with darkened sclerae and improved it for chimpanzee eyes with lightened sclerae. The benefit of white sclera in both observer species provides support for P1 beyond human observers alone. However, image size and brightness were varied together, so their separate effects could not be determined. The small sample of trained chimpanzees limits species-level generalisation.

Perea-García et al. (2026) extended the comparison to a live setting. Human observers judged the gaze of one model at two distances (1.3 and 5.3 m) and two illumination ranges (3–10 and 400–550 lux). The human-like condition retained the model’s natural light sclera and blue-green irises; the nonhuman-like condition used black scleral contact lenses surrounding the same irises. Human-like eyes had lower mean gaze-position estimation error in bright–near, bright–far, and dim–far conditions; nonhuman-like eyes had numerically lower mean error only in the dim–near condition, although this difference was not statistically supported. This descriptive pattern is consistent with a distance-related robustness advantage, but does not support a general human-like advantage under dim illumination. Its use of a live model preserves additional visual cues, but the single model, two levels of each viewing condition, and absence of a crossed observer–stimulus species design limit generalisation.

Together, these findings support P1 under some conditions while leaving the breadth of the advantage unclear. Ideally, tests would sample multiple, ecologically relevant levels of distance and illumination, including intermediate values rather than only two extremes. They should also distinguish ambient illumination from local shading and include peripheral viewing and brief observation. Crossed observer–stimulus species designs would further help assess observer-specific effects. Behavioural experiments are substantially more demanding than image analyses, however, particularly with nonhuman primates, and restricting the conditions tested is often unavoidable. Coverage can therefore accumulate across complementary experiments, with image analyses guiding the selection of informative conditions. Controlled images and live stimuli also offer complementary advantages: images permit precise manipulation, while live stimuli preserve motion, depth, and corneal reflections. Artificial eye animations demonstrate that corneal reflections can influence perceived gaze (Anstis, 2018), but their contribution across viewing conditions remains unclear. Targeted behavioural tests of these additional cues would help connect image-based predictions to the information observers actually use.

### Falsifiability and scope of P1

P1 would be falsified if comparisons across a varied, ecologically relevant range of viewing conditions ruled out a robustness advantage for human-like eyes. Relevant evidence would include equivalent or better maintenance of eye-gaze direction discrimination with pigmented sclerae as viewing conditions change. Conversely, an advantage confined to a narrow set of conditions would restrict the scope of P1.

## Prediction 2 Benefits for human group activities

P2 is that more robust eye-gaze direction discrimination enables finer adjustments of behaviour during group activities such as joint action, coordinated search, and mutual monitoring. Specifically, compared with sustained, close-range dyadic interaction, the benefits of human-like eyes are predicted to be greater when spatial dispersion creates variable viewing conditions or when monitoring more group members increases attentional demands.

The rationale for this prediction builds on the finding that observers accurately discriminated gaze direction from both human and chimpanzee eye images when the images were larger and brighter, whereas white-sclera eyes yielded higher accuracy when image size and brightness were reduced (Kano et al., 2022b). Spatially distributed group members may be farther away, shaded, or viewed peripherally. P1 concerns the discriminability of individual gaze cues; the added demand here is allocating limited attention across more individuals, analogous to multi-target tracking (Alvarez and Franconeri, 2007). Shorter viewing time per person overlaps with P1, and attention can itself alter perceptual sensitivity (Pestilli and Carrasco, 2005). The key question for P2 is therefore whether these perceptual benefits yield greater gains in monitoring and coordination under the viewing and attentional demands of group life, a context particularly relevant to the social functions of gaze.

### Social functions of gaze in interaction

P2 asks whether robust eye-gaze information improves coordination; studies of gaze’s social functions identify potential benefits and costs to test. Gaze contributes to gathering information about others, signalling attention and interpersonal attitudes, regulating conversational turns, and coordinating shared activities (Cook, 1977; Kleinke, 1986; Emery, 2000). Eye contact influences attention, face processing, and affective responses (Senju and Johnson, 2009; Hietanen, 2018), while joint attention and communicative cues support learning and coordinated action (Tomasello, 1995; Tomasello et al., 2007; Csibra and Gergely, 2009). Being observed can also influence prosocial behaviour, although its effects depend on context and should be distinguished from the less consistent effects of artificial watching-eye images on generosity (Bradley et al., 2018; Northover et al., 2017).

Much of this literature treats gaze broadly, combining eye direction, head orientation, and sometimes body orientation. Studies that isolate the contribution of visible eye-direction information during interaction are more directly relevant to P2, but are less common. Perea García et al. (2017) used goggles that concealed the eyes during a face-to-face cup-arrangement task. When eye information was available, dyads completed more configurations, showed a lower probability of conversational repair, and used visual–verbal strategies more frequently. Boucher et al. (2012) similarly found that concealing an informer’s eyes with sunglasses slowed the target-location phase of a verbally instructed cube-moving task. These experiments show that eye information can contribute to dyadic coordination alongside head orientation and speech, although they conceal the eyes rather than manipulate scleral colouration.

P2’s functional tests should also assess costs, because revealing gaze can expose privately valuable information. Studies displaying recorded gaze positions show that observers can infer another person’s choices, and that instructed concealment reduces this predictability (Foulsham and Lock, 2015). In competitive hide-and-seek games, such information improved opponents’ success when gaze was recorded non-strategically, whereas live players adapted their gaze and concealed their intentions (Hausfeld et al., 2021). In a behavioural experiment with a single chimpanzee dyad, an informed subordinate withheld gaze towards high-value food and, in another condition, lured a dominant towards a lower-value alternative, although gaze was inferred from head orientation (Hall et al., 2017). These studies demonstrate information leakage and suggest behavioural strategies to conceal gaze direction, but do not directly test the effects of scleral appearance. A direct test of these costs could vary scleral colouration in matched avatars during cooperative versus competitive resource search, controlling head cues and the signaller’s visual access. Outcomes should include the signaller’s resources lost, payoff, and search efficiency, alongside observers’ gains.

### Screen-based tests with multiple faces

Screen-based tasks test one aspect of P2: whether human-like eyes improve monitoring of others’ gaze as viewing conditions and the number of individuals vary. Increasing the number of faces can itself raise attentional demands. To distinguish the effects of viewing conditions and attentional demands, screen-based tasks can vary the number of faces or monitoring requirements under matched viewing conditions (such as distance, illuminance, retinal eccentricity, and viewing duration), and separately vary viewing conditions while keeping the number of faces and monitoring requirements constant. As described above, Yorzinski and Miller (2020) examined gaze detection in displays of multiple faces. Observers searched for a direct-gaze face among averted-gaze faces, or vice versa. In eight-face arrays, natural white sclera shortened the time to first target fixation relative to iris-matched sclera for both large and small upright faces. A further experiment compared four- and eight-face arrays to test how this time increased with the number of faces. For small upright faces, the increase was smaller with white than iris-matched sclera for both target directions; for large upright faces, this difference was not statistically significant. This pattern is consistent with P2’s prediction that white sclera can facilitate gaze monitoring as the number of faces increases. The pattern also suggests that this advantage depends on viewing conditions, such as face size, although the interaction tests did not provide statistical support for this possibility.

Sweeny and Whitney (2014) showed that observers can integrate gaze information across one to four faces to estimate a crowd’s average gaze direction. Their task did not manipulate scleral colouration, but could be extended to test how eye appearance affects estimates of a crowd’s average gaze direction as the number of faces, viewing conditions, and attentional demands vary.

### Testing P2 in live interaction

The live-interaction studies reviewed above show that eye information can contribute to dyadic coordination, but do not establish P2’s predicted context-dependent advantage. Testing this requires comparing sustained, close-range dyadic interaction with interactions where viewing conditions vary across space or monitoring more individuals raises attentional demands. Collective search, joint action, and mutual monitoring as relatively large groups offer relevant contexts: participants may use gaze to identify where others are searching, anticipate their next action, or determine who is attending to an event.

Restricting other communicative cues can help isolate the contribution of eye-direction information, whereas allowing them tests its additional value during multimodal interaction. When eye information is reduced, participants may compensate through more explicit speech, pointing, or head movements. Such compensation may preserve final task accuracy while increasing communication, coordination time, or effort. Alternatively, it may maintain performance without these additional costs. Comparisons should therefore assess both task outcomes and the adjustments needed to achieve them, and whether the benefits of human-like eyes increase when viewing conditions vary or attentional demands increase.

### Falsifiability and scope of P2

#### Screen-based tasks

P2’s monitoring prediction would be challenged if comparisons across a varied range of relevant conditions ruled out an advantage of human-like eyes in detecting or monitoring gaze, or an increase in that advantage as the number of faces or viewing difficulty increased. Benefits confined to a narrow set of conditions would limit the prediction’s scope.

#### Live tasks

P2’s functional prediction would be challenged if comparisons across relevant activities ruled out benefits of human-like eyes for coordination, task performance, or communication costs, or ruled out greater benefits under variable viewing conditions or higher attentional demands than during sustained, close-range dyadic interaction. A similar positive benefit across contexts would support a general functional role without supporting the predicted context-dependent advantage. Improved discrimination without functional gains, or compensation by other cues without additional time or effort, would weaken P2’s predicted functional benefit. Benefits need not occur in every activity, but effects confined to a very narrow set of contexts would weaken P2’s proposed relevance to human group life.

Information-leakage costs alone do not falsify P2, but would challenge P4’s selection scenario if evolutionary models ruled out the spread or maintenance of depigmentation across empirically plausible balances of costs and benefits.

## Prediction 3 Receiver sensitivity to eye-gaze cues

P3 is that, compared with other primates, humans show greater receiver sensitivity to eye-gaze cues across development. Here, receiver sensitivity includes both eye-gaze direction discrimination and the tendency to attend to and rely on eye-gaze cues (Table 1). Receiver sensitivity matters because directional information must be detected and used to influence interaction (Dawkins and Krebs, 1978). Like the CEH’s prediction of greater reliance on eye cues in humans (Tomasello et al., 2007), P3 arises from the possibility that signals and receiver sensitivity coevolved, rather than being a necessary premise of the gaze-signalling account. We first outline the comparative designs needed to test P3, then ask what developmental and task-specific studies reveal about its different components.

### Comparative studies for receiver sensitivity

To test P3, the most informative comparisons cross observer species with stimulus species, so that human and nonhuman observers encounter both human and nonhuman eye configurations. This helps distinguish receiver differences from differences in stimulus visibility or familiarity. Responses to head direction can additionally provide a reference within each observer because differences in motivation, training history, and stimulus familiarity can make absolute performance difficult to compare across species.

In Tomasello et al. (2007), human infants and nonhuman great apes followed a human demonstrator who looked upward using eye-only cues, head direction with closed eyes, or both together, with a neither-cue control. Human infants relied relatively more on eye direction, whereas apes relied relatively more on head orientation. By comparing responses to eye and head cues, the study identifies a difference in relative cue use, rather than merely in overall responsiveness, consistent with P3 and the cooperative eye hypothesis. The role of experience remains less clear. All apes were captive, and the three human-raised young chimpanzees showed no exceptional performance in this task, indicating that the pattern occurred despite experience with humans. However, experience was not quantitatively matched between groups, both groups observed only a human demonstrator, and developmental stages were not matched. Further comparisons crossing observer and stimulus species and accounting for experience and development are therefore needed to clarify the proposed difference in cue weighting.

Kano et al. (2022b) crossed human and chimpanzee observers with images of both species and demonstrated eye-gaze direction discrimination and benefits of white-sclera configurations in both observer species. Only three of ten chimpanzees completed the six-stage training, requiring several thousand formal training trials each. This training difficulty is tentatively consistent with P3, but it was an incidental finding rather than a standardised comparison between species. The study neither compared eye and head responses nor directly tested whether humans have greater receiver sensitivity to eye-gaze cues than chimpanzees. Training, motivation, general attention to eyes, and experience could all have contributed to the observed difficulty.

Deaner and Platt (2003) tested humans and rhesus macaques using eye-only and head-plus-eye cues. Eye-only cues produced cueing in both species, with no significant reduction relative to the combined cues in the reported analyses. This provides an important counterweight to a broad account in which strong responses to eye direction characterise humans but are generally weak in other primates. However, both observer species viewed macaque faces, there was no head-only condition, and the statistical analyses did not directly compare species.

### Early receiver sensitivity

To assess P3 across development, studies can compare when and how humans and other primates begin to respond to eye-gaze cues. Farroni et al. (2002) found that newborns preferred direct over averted gaze in frontal face photographs, and that direct gaze enhanced face-related processing in four-month-old infants. Farroni et al. (2004) found faster newborn orienting towards peripheral targets congruent with schematic pupil movements when those movements were visible. These findings demonstrate early preferences and orienting responses to eye information.

Comparable capacities have also been reported in nonhuman primates. Myowa-Yamakoshi et al. (2003) tested three infant chimpanzees, sampled between 10 and 32 weeks of age, with human faces differing in gaze direction. All three preferred direct over averted eyes in frontal faces when head orientation was held constant. In macaques, early mother–infant mutual gaze has been reported in naturalistic observations (Ferrari et al., 2009), although this does not isolate responses to eye direction from other aspects of face-to-face interaction. Neural processing of gaze has also been reported in humans (Emery, 2000; Itier and Batty, 2009; Senju and Johnson, 2009) and macaques (Perrett et al., 1985; Shepherd et al., 2009; Shepherd, 2010). Together, these studies establish overlapping gaze-related capacities in humans and other primates, while leaving quantitative differences in receiver sensitivity and its development unclear.

### Task-specific findings across behavioural paradigms

Studies of eye-cue use in other primates span object choice, gaze cueing, overt gaze following, and trained discrimination. These paradigms provide complementary measures of how observers detect and use eye-gaze information under different task demands.

### Experimenter-given cues in object-choice tasks

Object-choice tasks require observers to use an experimenter’s cues, such as pointing, touching or gaze direction, to select a rewarded object or location. These studies commonly compare head-plus-eye cues, in which both the head and eyes are directed towards the target, with eye-only cues, in which the head remains facing the observer while the eyes alone are directed towards the target. Observers must detect the cue and use it within the task’s referential or learned structure to succeed.

Itakura and Tanaka (1998) tested two chimpanzees and an orangutan across successive cue conditions and found successful use of both head-plus-eye and eye-only cues. The chimpanzees reached criterion in the minimum two sessions for each head-plus-eye condition, compared with three and seven sessions for eye-only cues. Peignot and Anderson (1999) found successful head-plus-eye cue use in four gorillas after training, but no successful eye-only use in the three tested with that cue. Agitation and nonresponses during eye-only trials limit interpretation of this contrast. Barth et al. (2005) found group-level use of head-plus-eye cues when chimpanzees entered the test area on each trial with the cue already present and maintained until choice, but not when they stayed there between trials and chose after cue presentation ended. Eye-only cues yielded no group-level success in either procedure, although one chimpanzee performed above chance. Two of three capuchins learned to use eye-only cues. When head and eye directions conflicted, both tested individuals favoured head orientation, significantly so in one individual (Vick and Anderson, 2000). Ten common marmosets inconsistently used combined head, eye and slight trunk cues in a two-container, single-choice task. In a nine-container task allowing two choices, the group used head-and-eye and subsequently eye-only cues following touch and pointing sessions, although eye-only success varied across sessions (Burkart and Heschl, 2006).

### Gaze-cueing tasks

A common gaze-cueing paradigm presents a nonpredictive gaze cue followed by a target either in the cued direction (congruent trials) or elsewhere (incongruent trials). Faster responses or higher perceptual accuracy on congruent than incongruent trials indicate that the gaze cue facilitated target processing. Tomonaga (2007) found no reliable reaction-time cueing from static eye-only cues in photographed or schematic human faces in two chimpanzees. Quasi-dynamic head-turn cues produced an effect under a modified procedure. However, fixation was not enforced and eye movements were not recorded. As described above, Deaner and Platt (2003) demonstrated eye-only gaze cueing in rhesus macaques, with shorter saccadic reaction times on congruent than incongruent trials. Shepherd et al. (2006) further showed that cueing by familiar conspecific faces depended on social status. Low-status observers made faster saccades to targets in the cued direction regardless of the depicted monkey’s status, whereas high-status observers showed this effect only for high-status faces. The stimuli included both eye-only and combined head-and-eye cues, but the paper did not report the status effect separately for the two cue types. Using a similar paradigm, Shafiei et al. (2026) measured detection accuracy for a near-threshold brightness increase in a peripheral light, with central fixation required until target onset. Head-plus-eye cues improved accuracy in both macaques, whereas standard eye-only cues did not produce significant cueing effects. An exploratory condition combining an enlarged avatar with increased eye displacement produced an eye-only effect in the one monkey tested.

### Overt gaze following

Following another individual’s head orientation has been reported across primates, other mammals, birds, and reptiles (Bugnyar et al., 2004; Bräuer et al., 2005; Kaminski et al., 2005; Shepherd, 2010; Zeiträg et al., 2023; Delacoux et al., 2025). Tomasello et al. (1998) found that four monkey species and chimpanzees followed a conspecific’s gaze towards a distant food item when attending to that individual. Bräuer et al. (2005) reported following of a human experimenter’s gaze in all four nonhuman great-ape species. Although neither study separated head and eye contributions, these findings provide a comparative baseline for examining eye-only following.

Several chimpanzee studies demonstrate eye-only following. Povinelli and Eddy (1996) found that five- to six-year-old chimpanzees turned to look above or behind themselves in response to a human’s eye-only cues, as well as to head-plus-eye cues. The two cue conditions did not differ significantly, although this did not establish precise directional discrimination. Okamoto et al. (2002) found that one infant chimpanzee reliably followed a human’s eye-only cues towards visible objects by 13 months. Follow-up tests of the same infant at 13–20 months also recorded eye-only following towards one of two identical toys positioned between the infant and the experimenter (Okamoto et al., 2004). Ferrari et al. (2000) tested pig-tailed macaques (*Macaca nemestrina*) with live head-plus-eye and eye-only cues from a human experimenter. At the group level, adults followed both cue types, whereas juveniles showed significant following relative to controls only for head-plus-eye cues. Individual analyses identified eye-only following in a five-year-old as well as in all adults.

### Comparative interpretation and further tests

Some findings suggest that eye-only cue use in nonhuman primates is more limited than responses to cues containing head orientation, consistent with P3. Yet this impression remains tentative because the studies are heterogeneous and behavioural responses can be influenced by training, motivation, general attention to eyes, and experience. Conversely, successful eye-cue use by particular individuals, in particular tasks, or in a single species establishes that the capacity is present under those conditions.

Testing P3 therefore requires crossed observer–stimulus species designs, in which human and nonhuman observers are tested with both human and nonhuman stimuli. Head-direction responses can provide a reference, while tasks and developmental stages should be made comparable to help distinguish receiver sensitivity from stimulus visibility and experience. Existing studies provide elements of this approach, but none of those reviewed here combines a crossed observer–stimulus species design with comparable tasks and developmental stages to test the proposed receiver specialisation.

### Falsifiability and scope of P3

P3 would be challenged by evidence of equivalent or greater receiver sensitivity in other primates across comparable developmental stages and viewing conditions in crossed designs. Discrimination, spontaneous cue use, and relative cue weighting should be assessed separately; comparing responses to eye and head cues helps distinguish differences in relative cue use from differences in overall responsiveness. Greater receiver sensitivity in humans, if confined to particular components, tasks, or developmental stages or inconsistent across response systems, would require the proposed specialisation to be narrowed.

## Prediction 4 Heritable variation and selection

P4 is that heritable variation in eye morphology was available when selection acted, and that the fitness benefits of eye-gaze signalling could exceed its costs for individuals bearing the relevant traits (Table 2). This prediction concerns the evolutionary link between the functional benefits considered under P1–P2 and changes in eye morphology. Direct evidence for this link remains limited: variation in appearance and present-day perceptual benefits do not by themselves establish selection mediated by eye-gaze signalling.

### Testing heritable variation and selection

Heritability can be investigated by combining standardised measurements of eye morphology with pedigree information or genomic estimates of relatedness. Repeated measurements would help distinguish stable differences among individuals from changes associated with age, disease and environment. The extent and uniformity of pigmentation on the exposed ocular surface could be measured using standardised imaging methods. Quantitative genetic models could then estimate additive genetic variance and heritability while accounting for shared environmental influences (Wilson et al., 2010).

Several sources of variation need to be distinguished when identifying suitable samples. Studies of the visible human scleral region have reported a darker, redder and yellower appearance at older ages (Russell et al., 2014) and population differences in superficial pigmentation of the overlying conjunctiva (Norn, 1985). Chimpanzees also show developmental changes, with many initially light-eyed infants acquiring pigmentation as they mature (Clark et al., 2023). Across primate species, associations between ocular colouration and latitude further suggest environmental influences (Perea-García et al., 2022, 2024). Such variation in scleral appearance does not necessarily imply differences in the visibility of eye-gaze direction as discussed above.

The occurrence of white sclera in adult chimpanzees such as within the Ngogo population provides a promising candidate for studying more pronounced within-population variation (Clark et al., 2023). Building on the longitudinal changes documented by Clark et al. (2023), family-based sampling could test whether persistent adult differences have a heritable component. Human ocular pigmentation is also variable (Perea-García et al., 2025). Individuals with marked pigmentation present from early life could provide additional candidates if this can be established as stable, nonpathological variation.

Functional benefits and costs could first be estimated in the human group activities considered under P2. Experiments could examine how eye appearance affects coordination and information leakage while other communicative cues remain available. Where opportunities arise in captivity, studies could examine interactions between white-sclera chimpanzees and conspecific receivers that have already demonstrated eye-only gaze following with human experimenters (Povinelli and Eddy, 1996; Okamoto et al., 2002, 2004). In wild populations such as Ngogo, studies could examine whether survival differs between individuals with white and dark sclera. These studies, however, would entail practical constraints: captive comparisons would require sufficient numbers of individuals with contrasting eye appearances and comparable backgrounds, while wild survival analyses would require long-term follow-up accounting for age, health, and social circumstances.

These practical constraints make evolutionary simulations a useful next step, using benefits and costs estimated from social interactions to examine the conditions under which depigmentation could evolve. For example, individuals could vary in a heritable eye-appearance trait that affects how reliably others obtain directional information under the viewing conditions examined in P1. The resulting information could improve coordination, as proposed in P2, but could also be exploited by competitors. Payoffs from these interactions could determine reproductive success in the model, allowing tests of whether a rare depigmented variant spreads and whether an established depigmented population resists invasion by more pigmented variants. Cooperative behaviour and eye appearance should be allowed to vary independently, rather than defining individuals with conspicuous eyes as cooperators.

Such models could vary the frequency of cooperative and competitive encounters, repeated interactions, partner choice, reputation, group structure, and receiver sensitivity. They could ask how large coordination benefits must be, how much information-leakage cost can be tolerated, and whether individuals with less readable eyes can benefit from others’ information without providing comparable information themselves. Repetition and reputation would be candidate mechanisms to test, rather than assumptions that automatically ensure stability. The aim is to identify the combinations of conditions under which depigmentation is favoured, including conditions under which it is not.

Existing models provide relevant starting points. Unakafov et al. (2020) showed that action visibility can enhance cooperation in an iterated coordination game but suppress it in an iterated Prisoner’s Dilemma. Mesoudi et al. (2025) modelled information sharing and free-riding, including how reputation-based partner choice can provide returns to information sharers. These studies concern action visibility and information sharing, respectively, rather than the evolution of eye morphology. Extending such approaches to an evolving eye-appearance trait would make the assumptions behind P4 explicit. Empirical findings should constrain the relevant benefits, costs, and perceptual parameters; a successful simulation would establish evolutionary feasibility under those assumptions, rather than demonstrate the historical occurrence of that process.

### Considering potential selective pressures on different aspects of eye morphology

Changes in scleral exposure, pigmentation, and depigmentation may reflect different selective pressures. To guide tests of P4 with the evidence currently available, we consider the possible communicative, biomechanical, and physiological functions of each change. The SGSH specifically concerns selection for gaze communication, so these comparisons help identify research priorities within that account.

Increased scleral exposure may have served communicative functions. Kobayashi and Hashiya (2011), for example, related scleral exposure to group size and neocortex ratio. However, noncommunicative explanations have also been proposed. Kobayashi and Kohshima (2001) reported an association between body size and scleral exposure and proposed that larger eye openings may facilitate visual scanning through ocular rotation, reducing reliance on head reorientation. Both social and biomechanical explanations for scleral exposure are therefore plausible.

Pigmentation of exposed ocular surfaces could serve photoprotective or other physiological functions. Comparative associations between eye colouration and environmental conditions are relevant to this possibility (Perea-García et al., 2022, 2024). White scleral regions have been documented in nonhuman primates (Clark et al., 2023; Silva et al., 2017), and light scleral appearance also occurs in other mammals (Caspar et al., 2023). However, white regions may be concealed rather than externally exposed (Silva et al., 2017). For the SGSH, what requires explanation is therefore why externally exposed sclera is depigmented rather than pigmented, not the presence of white scleral tissue itself.

The gaze-camouflaging hypothesis proposes that pigmentation could reduce the eye-direction information available to competitors or other observers (Kobayashi and Kohshima, 1997, 2001). Chimpanzees and mountain gorillas can combine dark sclera with relatively light irises, retaining conspicuous contrasts within the face (Perea-García et al., 2019; Kano et al., 2022a). Receiver-vision models likewise show detectable regional contrasts in chimpanzee eyes at ecologically relevant distances (Whitham et al., 2022a). Pigmentation therefore does not necessarily make the eyes inconspicuous; a camouflage benefit beyond close range would require pigmentation to reduce useful eye-direction information not already provided by visible head-direction cues, and thereby reduce information-leakage costs. As reviewed under P3, eye-only cue use varies across species and contexts, so such a benefit cannot be assumed from eye colouration alone. Testing this possibility therefore requires separating visibility of eye-gaze direction, eye-gaze direction discrimination, and eye/head-direction use, and measuring whether pigmentation actually lowers information-leakage costs for the observed individual.

### Alternative origins and functions of scleral depigmentation

The P1-3 findings provide a functional basis for testing whether depigmentation of exposed sclera was favoured through gaze communication. Assessing this possibility under P4 also requires considering how depigmentation could have arisen through other processes and what other functions it may serve. The following section examines these alternatives.

Under the self-domestication hypothesis, reduced scleral pigmentation may have arisen as a by-product of selection against aggression, with subsequent selection potentially acting directly on any heritable variation in this trait (Hare, 2017). However, the absence of a consistent association between domestication and scleral appearance across mammals limits the generality of this explanation (Caspar et al., 2023). Ecological and physiological processes, including those considered above, also remain relevant alternatives. A configuration arising through these processes could later be maintained or modified by its effects on eye-gaze signalling.

Scleral appearance also conveys social information other than eye-gaze direction. Experimental changes in scleral colouration influence perceived attractiveness and emotional state (Provine et al., 2013a, 2013b), as well as trustworthiness (Wacewicz et al., 2022). Typically developing children and adults prefer stuffed animals with visible white sclera, whereas children with autism spectrum disorder in the same study showed no consistent preference (Segal et al., 2016). Five-year-old children also show preferences for faces with white rather than dark sclera as potential cooperative partners (Wolf et al., 2023). Perea-García et al. (2025) discuss broader explanations based on health, sexual selection, and non-referential social signals. These functions can coexist with eye-gaze direction signalling. Evidence for them therefore does not by itself contradict the proposed directional-signalling function; their respective contributions to the evolution of depigmentation remain to be established.

### Falsifiability and scope of P4

The proposed selective mechanism would be weakened if functional and fitness-related studies ruled out a net benefit of eye-gaze signalling associated with the relevant morphology under the conditions specified by the account. Evolutionary simulations would further challenge a particular scenario if depigmented variants failed to spread when rare or could not persist once established across the empirically supported conditions proposed to favour them. A scenario requiring implausibly large coordination benefits, negligible information-leakage costs, or unsupported assumptions about repeated interaction or partner choice would also be weakened.

Heritability and historical selection require separate interpretation. Evidence ruling out appreciable additive genetic variation today would constrain an ongoing evolutionary response to selection, but would not exclude variation available during past selection. Conversely, finding heritable variation would not demonstrate that gaze signalling drove its evolution. Evidence that the trait originated through another process would challenge a gaze-signalling account of its origin while leaving possible later maintenance through signalling benefits open to testing. P4 therefore requires explicit claims about the trait, evolutionary stage, and selective conditions being explained.

## Future directions summary

The main gaps in current knowledge differ across the four predictions. For P1, image-based measures and behavioural performance need to be linked across the variation in eye appearance and viewing conditions encountered in everyday interactions. For P2, screen-based tasks should test whether human-like eyes improve gaze monitoring as the number of faces or viewing difficulty increases. Live tasks should test whether these eyes provide greater coordination benefits under variable viewing conditions or higher attentional demands than during sustained, close-range dyadic interaction, and whether other communicative cues compensate without additional time or effort. For P3, crossed observer–stimulus species designs are needed to compare receiver sensitivity across development using separate measures of perceptual discrimination, spontaneous eye-cue use, and relative eye–head cue weighting, with responses to head direction as a comparative reference and controls for training, motivation, general attention to eyes, and experience. For P4, studies of heritable morphological variation need to be complemented by evolutionary simulations examining when the benefits and costs estimated from functional studies permit depigmentation of exposed sclera to spread and persist. Together, the tests in Table 2 connect morphology, perception, interaction, and evolution while allowing each prediction to be evaluated separately.

## Conclusion

Head or body orientation can indicate gaze direction when gaze shifts follow those movements, whereas eye-direction cues become informative when ocular mobility allows gaze to diverge from head or body orientation (Emery, 2000). Scleral exposure may facilitate visual scanning through ocular rotation (Kobayashi and Kohshima, 2001), while pigmentation of exposed ocular surfaces may provide photoprotection (Perea-García et al., 2024); neither explanation requires a communicative origin. Scleral depigmentation producing a human-like white appearance has been observed in some wild chimpanzees (Clark et al., 2023). Sensitivity to eye-gaze cues has been observed in some individuals of nonhuman ape and monkey species (Deaner and Platt, 2003; Ferrari et al., 2000; Povinelli and Eddy, 1996). Pigmented nonhuman ape eyes already permit eye-gaze direction discrimination when clearly visible at close range (Kano, 2023). The presence of these traits and capacities alone is therefore not what the SGSH needs to explain. Its specific question is whether depigmentation of exposed sclera provides greater social benefits under variable viewing conditions or higher attentional demands than during sustained, close-range dyadic interaction, and whether these benefits can explain the trait’s spread and maintenance in humans even if it initially arose through other processes.

To answer this question, the SGSH links four predictions concerning perceptual robustness, context-dependent coordination benefits, comparative receiver sensitivity, and the heritable variation and selective conditions required for evolutionary change. Comparisons between humans and other great apes are particularly informative because scleral exposure occurs in both, while variation within and among species provides further tests of the morphological and behavioural predictions. Existing findings support perceptual benefits under some conditions, while evidence for the other predictions remains more limited. Explicit tests and falsification criteria can help guide future research on the SGSH. Addressing these questions would help resolve a fundamental question in human evolution: whether humans evolved morphological traits for conspecific communication, specifically to monitor one another’s attentional foci.

## Acknowledgements

The authors thank Dr J. O. Perea-García and Dr S. Wacewicz for valuable discussions, and two anonymous reviewers for constructive comments that helped improve this review.

## Author Contributions

F.K. conceived the review, developed the Stabilized Gaze Signalling Hypothesis (SGSH), conducted the analyses, and wrote the original draft of the manuscript. H.K. and K.H. contributed to manuscript revision and editing. All authors reviewed and approved the final manuscript.

## Research Transparency and Reproducibility

The analyses reported in this article were conducted using previously published data. Details of data sources, data processing, and analytical procedures are provided in the Supplemental Material.

## Data Availability

Data and code supporting the analyses are available via Open Science Framework: https://osf.io/z6753/

## Declaration of generative AI and AI-assisted technologies in the writing process

During the preparation of this manuscript, the authors used ChatGPT 5 and 6 to improve language and readability. Following the use of this tool, the authors carefully reviewed and edited the content and take full responsibility for the content of the publication.

