## Supplemental Material for "The stabilized gaze signalling hypothesis and the evolution of human eye appearance"

### Materials and Methods

This reanalysis incorporated the dataset used in Kano et al. (2022a) and followed the same analytical protocol. Codes, data, and image thumbnails are available at Open Science Framework (https://osf.io/z6753/?view_only=5f6f393077194fb789c7fe5c44bbe06e).

**Samples.** A detailed summary of the samples in Kano et al. (2022a) is provided in Table 1 of that paper. Briefly, the dataset comprised 160 high-resolution facial images representing seven great-ape species—bonobos (*Pan paniscus*), chimpanzees (*Pan troglodytes*), mountain gorillas (*Gorilla beringei beringei*), western lowland gorillas (*Gorilla gorilla gorilla*), Bornean orangutans (*Pongo pygmaeus*), Sumatran orangutans (*Pongo abelii*), and humans (*Homo sapiens*). Human photographs were drawn from the Columbia Gaze Dataset (Smith et al., 2013), which includes individuals of diverse ethnic backgrounds.

For comparison, 20 additional eye images were extracted from the high-resolution version of Figure 4 in Perea-García et al. (2025). Each image (123 × 73 pixels) corresponds to the approximate eye-region size used in Kano et al. (2022a). Five of these images showed averted eyes and were classified as “averted,” whereas the remaining fifteen were “direct.” The shape analysis uses the direct-eye set only; the color analysis included both categories. For the shape analysis, ten publicly available, forward-facing eye images each were added for the golden langur (*Trachypithecus geei*) and the southern pig-tailed macaque (*Macaca nemestrina*). All new images were oriented toward the camera and showed neutral expressions (see Fig. 2A for examples). For the color analysis, following Kano et al. (2022a), overall brightness and contrast of the images (including both eye and skin regions) were standardized in Photoshop by automatic level adjustment (0–255 range, gamma correction to center the histogram around mid-gray).

**Shape analysis.** Eye contours and iris borders were manually traced (2-pixel width) in Photoshop and converted to binary masks in MATLAB (R2024b, MathWorks). Two standard indices were then calculated: 1) Sclera Area Index, the ratio of scleral area to total eye-opening area (a two-dimensional measure; Kano et al., 2022a). 2) Sclera Size Index, the ratio of the longest eye-opening diameter to the longest iris diameter (a one-dimensional measure; Kobayashi & Kohshima, 2001). These indices quantify relative scleral exposure independently of overall eye size.

**Color analysis.** The analysis focused on how clearly the iris and sclera boundaries stood out from adjacent facial regions across the six schematic gaze-visibility patterns illustrated in Figure 3A. Color contrasts between features were calculated in the CIE LAB color space, which approximates trichromatic primate vision. The color difference between any two regions was expressed as

$$\Delta E = \sqrt{(L_1 - L_2)^2 + (a_1 - a_2)^2 + (b_1 - b_2)^2},$$

where $L$, $a$, and $b$ represent lightness and the red–green and blue–yellow axes.

Separate filled and outline masks were prepared for the iris, sclera, eye outline, and surrounding skin. To capture boundary regions, each 2-pixel tracing was dilated to an 8-pixel outline (i.e., 3 pixels expansion on either side). Masks were mutually exclusive; pupils and specular reflections were excluded. Mean LAB values were calculated for each mask, and pairwise ΔE values were obtained to quantify contrast. This procedure yields size-independent measures of eye-feature conspicuousness.
